# Characterizing the impact and maintenance of specialized toxin tolerance in a generalist species

**DOI:** 10.1101/2023.08.24.554702

**Authors:** Grace Kropelin, Clare H. Scott Chialvo

**Affiliations:** Department of Biology, Appalachian State University, Boone, NC

**Keywords:** Plant-Insect Interactions, Host Preference, Competition, *Drosophila guttifera*, Cyclopeptides, *Amanita*

## Abstract

Understanding how plant-herbivore interactions can drive coevolution is a central goal of ecology and evolutionary biology. Of particular interest are the defenses produced by host plants/fungi and their impact on herbivore feeding strategies. In the *immigrans-tripunctata* radiation of *Drosophila*, mushroom-feeding species are classified as generalists, but their acceptable hosts include deadly *Amanita* species. In this study, we used behavioral assays to assess whether the mushroom-feeding species *Drosophila guttifera* is becoming a specialist and to characterize the impact of competition on host usage. We conducted feeding assays to confirm the presence of cyclopeptide toxin tolerance. We then completed host preference assays in female flies and larvae and did not find a preference for toxic mushrooms in either. Finally, we assessed the effect of competition on oviposition preference. We found that the presence of a competitor’s eggs on the preferred host was associated with the flies increasing the number of eggs laid on the toxic mushrooms. Our results suggest that an adaptation associated with specialized feeding behavior is not altering host usage and that competition plays a role in maintaining this trait. More broadly our work highlights how access to a low competition host resource helps to maintain adaptations with fitness costs.

## INTRODUCTION

Understanding the interactions between plants and their herbivores and how they can drive coevolution is an active area of research. While some associations can be mutualistic in specific instances (de Mazancourt et al. 2001; Johnson et al. 2021), the primary focus is on the antagonistic interactions where the herbivore is classified as a parasite of its host (Berenbaum and Feeny 1981; Ehrlich and Raven 1964; López-Goldar et al. 2022). In plants and fungi, interactions with herbivores lead to the evolution of a variety of defense mechanisms that include mechanical structures, chemical compounds, and phenological shifts (Agrawal et al. 2009; Fraenkel 1959; Hanley et al. 2007). These defenses, in turn, place pressures on herbivores and select for adaptations that allow them to avoid or mitigate the negative costs (Becerra 1997; Futuyma and Moreno 1988).

Of particular interest is how the type of chemical defense influences the breadth of host usage in herbivores. Specifically, hosts that produce highly noxious compounds are expected to exclude generalist feeders that lack the necessary adaptations (Ehrlich and Raven 1964; Krieger et al. 1971). These generalist species instead possess tolerance mechanisms that are effective against less toxic host secondary metabolites that are found across many of their acceptable hosts. Conversely, hosts that produce acutely toxic defensive compounds are consumed by specialists which evolved tolerance, but lost the ability to feed on a wide variety of hosts (Cornell and Hawkins 2003; Whittaker and Feeny 1971). Despite these predictions, there are cases where generalist herbivores evolve mechanisms that allow them to feed on heavily defended hosts (Dussourd and Denno 1994; Hartmann et al. 2005).

We know far less about how adaptations associated with tolerance to highly noxious host compounds evolve in, and impact the evolution of, generalist feeders. Given that these compounds would not be present in all or even most of a generalist’s acceptable hosts, tolerance mechanisms would experience inconsistent selection and might also have associated fitness costs. However, the ability to feed on a highly toxic host offers a potential escape from competition (Harrison and Karban 1986; Viswanathan et al. 2005; Zytynska and Preziosi 2013) and access to enemy free space (Atsatt 1981; Denno et al. 1990; Mulatu et al. 2004). One group of generalist feeders that possesses tolerance to highly toxic compounds is mushroom-feeding *Drosophila* in the *immigrans-tripunctata* radiation (reviewed in Scott Chialvo and Werner 2018). These flies are classified as generalist feeders of fleshy-white Basidiomycota, but can also feed on *Amanita* mushrooms that produce deadly cyclopeptides. While the mushroom-feeding flies can develop at natural concentrations of cyclopeptides (Jaenike 1985), this adaptation is rapidly lost (∼1 million years) when species transition to feeding on hosts other than mushrooms (Spicer and Jaenike 1996). While this suggests that there are fitness costs associated with maintaining tolerance in these flies, the novel adaptation is potentially impacting the life history of these species.

Novel adaptations, such as tolerance to highly toxic compounds, can initiate adaptive radiations and allow species to enter new ecological niches that release them from competition and offer enemy-free space. Among the mushroom-feeding *Drosophila* that are cyclopeptide tolerant, several species occur in the *quinaria* species group, a young adaptive radiation (∼19.5 million years; Izumitani et al. 2016). In this group, mushroom-feeding and cyclopeptide toxin tolerance are ancestral traits (Erlenbach et al. 2023; Scott Chialvo et al. 2019; Spicer and Jaenike 1996). As a result, we hypothesized that mushroom-feeding species in the *quinaria* group could be transitioning from a generalist to a specialist feeding habit. In this study, we use a variety of behavioral to assess host preferences in *Drosophila guttifera* and quantify whether there is any evidence of preference for toxic host mushrooms over those that are edible. This species is a member of the *quinaria* group, and some lines show a higher probability of survival on diets with cyclopeptides than without (personal observation). We also assessed the role of competition (intra- and interspecific) in host selection. In sum, our results suggest that female flies and larvae do not exhibit a preference for toxic host mushrooms, but the ability to use a host that reduces competition may act as a selection pressure on this trait.

## MATERIALS AND METHODS

### FLY STOCKS

For this study, we worked with two, unique genotypes of *Drosophila guttifera* (KD – 15130-1971.00; TW – 15130-1971.10). These two lines were originally maintained in the *Drosophila* species stock center; however, they are no longer available and were provided by Kelly Dyer (University of Georgia; 15130-1971.00) and Thomas Werner (Michigan Technological University; 15120-1971.10). The fly stocks were maintained on a standard diet consisting of Carolina 4-24 *Drosophila* instant medium supplemented with a piece of fresh, white mushroom (*Agaricus bisporus*). A dental cotton roll was placed in each vial as a substrate for pupation. The stocks and experiments were maintained at 23° C with a 12:12 light-dark cycle for the toxin tolerance and oviposition assays. Due to an incubator failure, the fly colonies and experiments for the larval preference and competition assays were maintained at room temperature with a 12:12 light-dark cycle.

### TOXIN TOLERANCE ASSAY

To quantify tolerance to an individual toxin and a complex mix in both *D. guttifera* genotypes, we conducted feeding assays in 7.5 mL glass scintillation vials containing 250 mg of an instant *Drosophila* medium (73.5%) and freeze-dried portabella mushrooms (*Agaricus bisporus*; 26.5%) mixture. The dried mix was resuspended using 1 mL of water or a solution containing the toxin(s): α-amanitin (250 µL α-amanitin and 750 µL deionized water) and natural toxin mix (185 µL toxin mix and 815 µL deionized water). The final concentration of the α- amanitin diet was 250 µg/g, which is equivalent to the mean concentration of this compound in *Amanita bisporigera* (Tyler et al. 1966; Yocum and Simons 1977). The concentration of the natural toxin mix was 100 µg/g. We found that toxin susceptible species do not survive on diets containing 50 µg/g (unpublished data).

We extracted the natural toxin mix from dried *Amanita phalloides* that were collected along Point Reyes, CA in December 2017. As cyclopeptides are thermostable (Li and Oberlies 2005), it is not expected that drying the mushrooms would affect the potency of toxins. The extraction was performed using an accelerated solvent extractor and two solvents, methanol:water (5:4 v/v) and methanol following the protocol described in Scott Chialvo et al. (2020). We used these two solvents because *Amanita phalloides* contains a mixture of 14 known cyclopeptide toxins with differing polarities (Faulstich et al. 1975; Munekata et al. 1978; Wieland 1968; Wieland 1983). The extracted solution was dried down in a rotary evaporator. As the phallotoxin subclass of cyclopeptides are less polar than amatoxins, the toxin concentrate was resuspended in a mixture of methanol and deionized water (325 mL and 550 mL respectively). We confirmed the concentration of amatoxins, the only cyclopeptide class to readily pass through the gut lining (Diaz 2005; Li and Oberlies 2005), using HPLC analysis and commercially available chemical standards (*i.e.*, α- and β-amanitin). The solution contained 0.541 µg/µL amatoxins (0.215 µg/mL α-amanitin and 0.326 µg/mLβ-amanitin). To account for the potential impact of methanol on survival, we added 56 µL of methanol to both the control and α-amanitin treatments. After adding the solutions to the instant food-mushroom mix, the scintillation vials were placed uncovered in a fume hood for 96 hours to allow the methanol to evaporate. Because some water could have evaporated, we then added 200 µL of DI water to each of the food vials.

Prior to adding fly larvae, a small piece of watercolor paper was placed into each vial for pupation. For each scintillation vial, we picked and placed 15 early, first instar larvae into each environment. For 30 days, the vials were monitored daily for survival to adulthood, which was based on successful emergence of adult flies from their pupal case. For both genotypes, we completed 5 replicates for each of the three dietary treatments.

### OVIPOSITION PREFERENCE

To characterize oviposition preference for both *D. guttifera* genotypes, we quantified the number of eggs laid on each of the three media: edible mushroom, toxic mushroom, and tomato. We used the tomato-based medium as a negative control because *D. guttifera* does not naturally use fruits, such as tomatoes, as hosts (Sturtevant 1921). The mushroom media consisted of 200 mL deionized water, 8.3 g dehydrated and ground up mushroom, 4 g fly agar, and 200 mg tegosept (an antifungal agent). The edible medium used dried portabella mushrooms (*Agaricus bisporus*) whereas the toxic medium used dried deathcaps (*Amanita phalloides*). The tomato medium was made with a 1:1 ratio of tomato juice to deionized water (100 mL each), 4 g fly agar, and 200 mg tegosept. These recipes are based on media used in oviposition assays of other fly species in the *immigrans-tripunctata* radiation (Jaenike 1986). The media were allowed to set in 15 mL falcon tubes.

Oviposition assays were conducted in sterile petri dishes (94 mm × 16 mm). The media were sliced into 5 mm thick disks that were placed in equal distance from each other. A mated, female *D. guttifera* (7 to 10 days after emerging from pupation) was placed in the center of the plate (Figure 1). The assays were kept in a 22.5°C incubator with a 12:12 light dark cycle for a total of 72 hours. The female fly was then removed and the number of eggs present on each medium was counted. We completed at least 50 complete replicates for each for both genotypes. A replicate was only considered successful if three or more eggs were laid.

**Figure 1.**
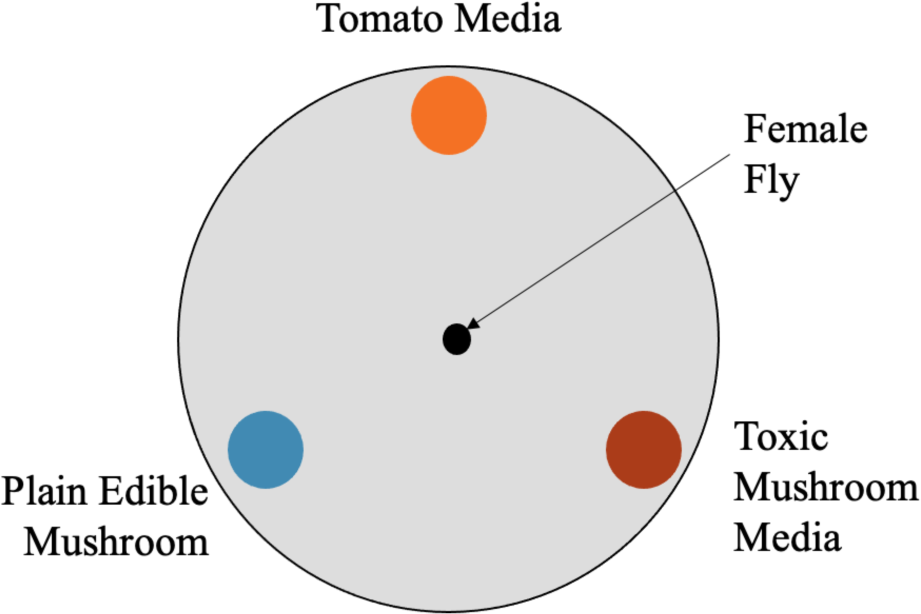
Organization of host media and female fly placement on petri dish for oviposition preference and competition assays.

### LARVAL PREFERENCE ASSAY

To characterize larval feeding preference for either toxic or edible mushrooms, we quantified the change in mass of three different media: plain agar, edible mushroom, and toxic mushroom. Due to the availability of dried *A. phalloides* and the results of the oviposition assay, we limited the larval feeding preference assay to the KD *D. guttifera* line. Plain agar medium was made with 200 mL water, 4 g fly agar, and 200 mg tegosept. The edible and toxic mushroom media used the same components as described above in the oviposition assays. Each of the three media were set using silicone square ice cube trays that produce 1” × 1” cubes. To these trays, we added 8mL of liquid media in each cube.

A cube of plain medium was placed in a small petri dish (35 mm × 10 mm). The dish was slotted into a 6 oz square bottom Drosophila bottle (Genesee Scientific) containing ≥50 adult *D. guttifera* KD flies (a mix of males and females) that were 7-10 days post emerging from their pupal case and are expected to be sexually mature. This setup was maintained at room temperature for 14 hours to allow for oviposition to occur. The number of eggs laid on each cube were counted, and we cut the cubes down the center so that each half had approximately equal numbers of eggs. A cube was not used if less than 10 eggs were present on each half. After the cube was cut in half, the mass was taken prior to placing onto a large petri dish (94 mm × 16 mm) for the larval preference assay.

For the preference assay, a half cube of plain medium was placed between a half cube of each mushroom medium. The cubes were arranged so as not to be in direct contact and separated by approximately 3 mm. As the majority of the eggs were found on the top of the cube and to minimize potential bias related to the placement of the cubes, one half of the plain agar medium was oriented toward the edible mushroom medium, and the other half was oriented toward the toxic mushroom medium (Figure 2). Once all the plates were labeled, we took the mass (mg) of both mushroom media and placed them on the plates. The plates were maintained at room temperature (22 °C) for 96 hours to allow the eggs to hatch and the resulting larvae to feed. Under these environmental conditions and this length of time, *D. guttifera* larvae reach late third instar just prior to pupation (personal observation). After 96 hours, we recorded the mass of each media again.

**Figure 2.**
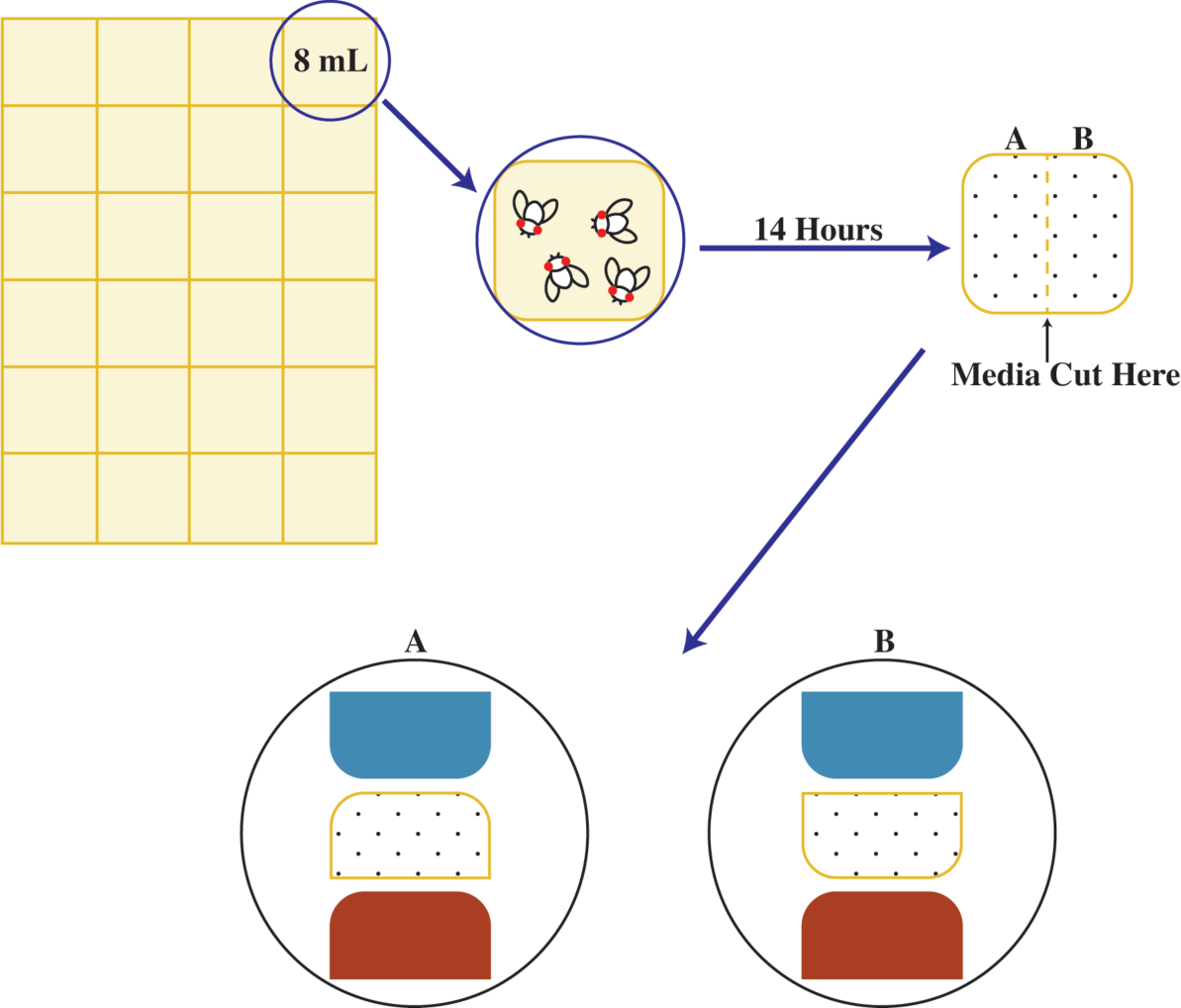
Experimental setup used in the larval preference assay. Flies laid on plain agar media over 14 hours. The media containing eggs was split in half and then situated between blocks of the edible and toxic mushroom media.

### COMPETITION ASSAY

To characterize the effects of competition on oviposition preference in the *D. guttifera* KD line, we measured how inter- and intraspecific competition impact the number of eggs laid and the location in comparison to no competition. The interspecific competitor used was *D*. *tripunctata* (2007 iso 1 Athens, GA line; provided by Kelly Dyer). Both *D. guttifera* and *D. tripunctata* occur in eastern North America from Texas to Florida and north into Canada (Werner et al. 2020). Thus, we expect these species would compete for resources in their natural distributions. For the intraspecific competition, we used eggs laid by females from the TW *D. guttifera* line.

The competition assays included the same media types made using the same protocols that we used in the oviposition assay (*i.e.*, edible mushrooms, toxic mushroom, and tomato (negative control)). For this assay, we allowed the three media to set in small petri dishes (35 mm × 10 mm). These petri dishes were filled to a depth of 5 mm.

For the assays, we compared the number of eggs laid in the presence of no competition and inter- and intraspecific competition. A replicate included each of these treatments run concurrently using female *D. guttifera* KD from the same brood. The competitors laid their eggs on the edible mushroom medium. We slotted the petri dish containing edible medium into a 6 oz square bottom Drosophila bottle containing ≥50 adults (mix of males and females; 7-10 days post emergence) of either *D. guttifera* TW or *D. tripunctata*. After 14 hours, we removed the dishes and checked for eggs.

We conducted the assays in a large petri dish (94 mm × 16 mm) filled with plain agar to a depth of 7 mm. The plain agar was made using the same protocol as in the larval preference assay. We used a 9 mm diameter leather awl to punch holes that were equidistant for the oviposition media (Figure 1). A 7 mm diameter hole was placed in the center of the dish for the female fly. To limit the access of *D. guttifera* KD flies to the top surface of the oviposition media, we placed the disks of the three media (9 mm wide) into equivalently sized holes in the plain agar. For the inter- and intraspecific competition treatment, the edible medium contained ≥4 eggs. The no competition treatment used a disk of edible medium without eggs or exposure to other flies. We placed a *D. guttifera* KD female into the central hole and maintained the dish at room temperature for 72 hours. After 72 hours, we removed the flies and documented the number of eggs laid as well as the locations (edible, toxic, tomato, plain, and near each oviposition medium). Eggs laid in the 7 mm hole that the fly was placed in were coded as being laid in plain medium. Eggs laid within 5 mm of an oviposition medium were coded as being near to that medium. We completed 50 replicates; replicates were only used if ≥ 3 eggs were laid in two of the treatments.

### STATISTICAL ANALYSES

All collected data (*i.e.*, survival to adulthood, number of eggs laid, and change in mass) were analyzed using JMP Pro v 16.0.0 (SAS Institute Inc.). For the toxin tolerance assay, we coded the survival of each individual larvae in a vial using a binary strategy (0 = deceased, 1 = adult successfully emerged). To assess the contribution of genotype (line), treatment (diet), and the interaction between the two factors on the probability of survival, we performed a logistic regression analysis using a generalized linear model with binomial distribution, logit link, and Firth bias-adjusted estimates (Firth 1993). Additionally, we also calculated a chi-square for the effect of treatment in each genotype to determine whether survival was significantly affected by the toxins.

For the other assays (oviposition preference, larval preference, and competition), we assessed the contribution of the factors of interest with a regression analysis implemented using a standard least square model and an effect leverage emphasis. For the oviposition preference assay, we assessed how genotype (line), oviposition media, and the interaction between these factors influenced the variation in the number of eggs laid. We also conducted a one-way ANOVA of the number of eggs laid on each media type (edible mushroom, toxic mushroom, and tomato) for each genotype to examine their preference for each of the medias. We compared the mean number of eggs laid on each media using a Tukey HSD analysis. In the larval preference assay, we assessed how the number of eggs on the plain media, media type, and the interaction between these two factors affected the variation in the change in mass. A one-way ANOVA of change in mass by media and Tukey HSD were used to compare the mean change in mass of each media type. For the competition assay, we first conducted a one-way ANOVA with Tukey HSD to assess how this factor influenced the total number of eggs laid. We used the regression analysis to assess the impact of competition, media, and their interaction on the variation where eggs were laid. To examine how oviposition preference varied under each competition treatment, we completed a one-way ANOVA of eggs laid by media with a Tukey HSD.

## RESULTS

To better understand the impact of a novel adaptation, cyclopeptide tolerance, on the life history of *D. guttifera*, we combined feeding assays with characterizations of both adult and larval host preference tests. Our goal was to determine whether this trait is inducing changes in feeding strategies and under what conditions is this trait adaptive.

### TOXIN TOLERANCE

To confirm that both available *D. guttifera* lines possess the trait of interest, cyclopeptide tolerance, we reared larvae on diets with and without a natural concentration of α-amanitin (250 µg/g) or a mix of cyclopeptide toxins (100 µg/g amatoxins) and measured survival to adulthood. While the probability of survival in both lines on each of the three diets was greater than 0.1, the responses of the two lines differed on the diet containing the single toxin (Figure 3). For the KD line, the probability of survival was highest on this diet (α-amanitin), but the TW line showed a decrease in comparison to the no toxin diet. In both lines, developing on a diet containing the toxin mix reduced the probability of survival in comparison to the no toxin diet. Although both lines showed differences in the probability of survival across the three diets, they were only significant in the TW line (*P <* 0.0001; Supplemental Table 1). In our examination of the variation in probability of survival across both *D. guttifera* genotypes, the genetic line accounted for the greatest contribution (*P =* 0.00002; Table 1). Line by diet interactions (*P =* 0.0277) and dietary treatments (*P =* 0.0009) also contributed significantly to the variation in this phenotype.

**Figure 3.**
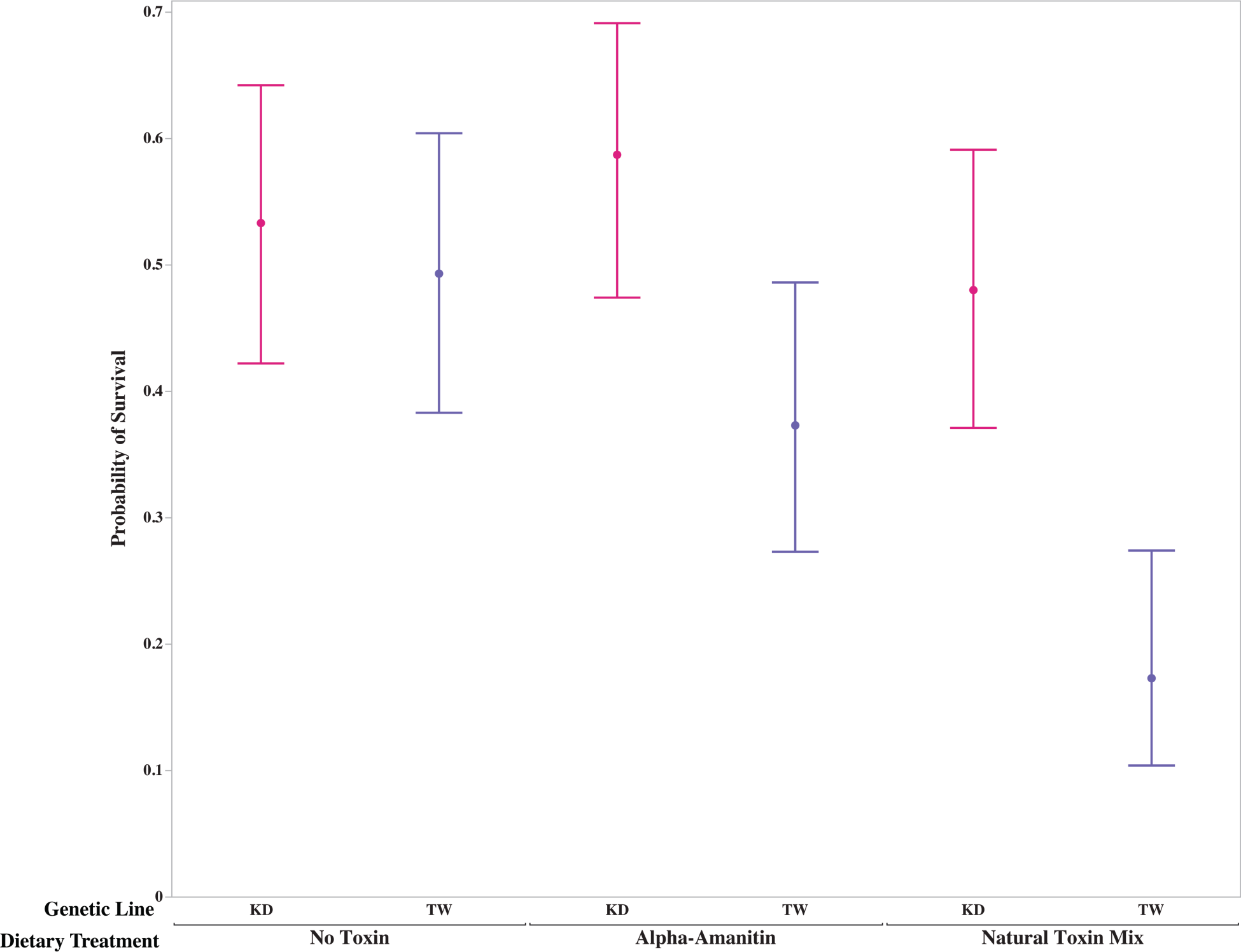
Comparisons of the line by diet interactions in the probability of survival to adult in both *D. guttifera* lines across the three dietary treatments. X-axis indicates the dietary treatment. The 95% confidence interval is included for the probability of survival in each genotype by treatment combination.

**Table 1.**
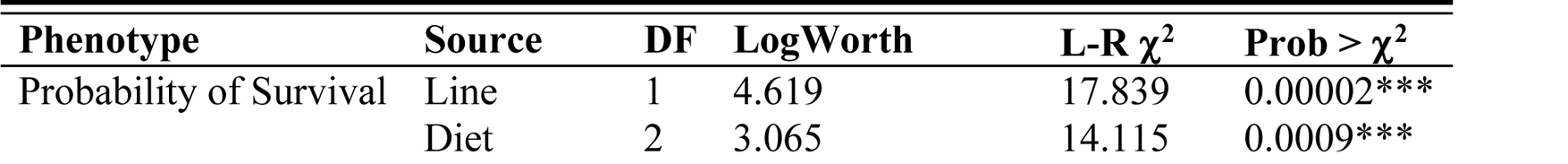
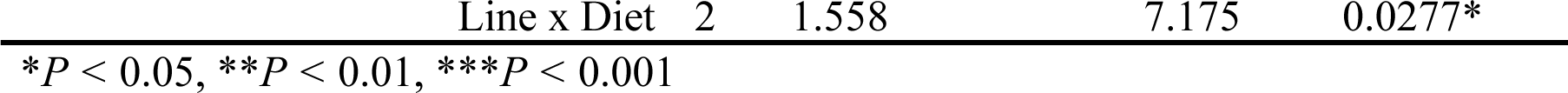
Analysis of variance in larval performance on diets with and without cyclopeptide toxins.

### OVIPOSITION PREFERENCE

To assess whether the female flies in both *D. guttifera* lines are partial to laying eggs on toxic mushroom hosts, we documented the number of eggs laid and the location when the flies were provided three host options (edible mushroom, toxic mushroom, and a negative control – tomato). Both genotypes laid the most eggs on the edible mushroom medium (mean number of eggs laid > 10; Figure 4). This was significantly higher than the mean number of eggs laid on either the toxic mushroom or tomato (negative control) media (*P <* 0.001; Table 2). Somewhat surprisingly, the flies laid slightly more eggs on the tomato-based medium (negative control) than the toxic mushroom medium. However, this difference was not significant (*P >* 0.05). When examining the sources of variation in this data, only the oviposition media type made a significant contribution (*P <* 0.0001; Table 3). Genetic line and the interaction between it and media did not contribute significantly (*P =* 0.273 and *P =* 0.418 respectively).

**Figure 4.**
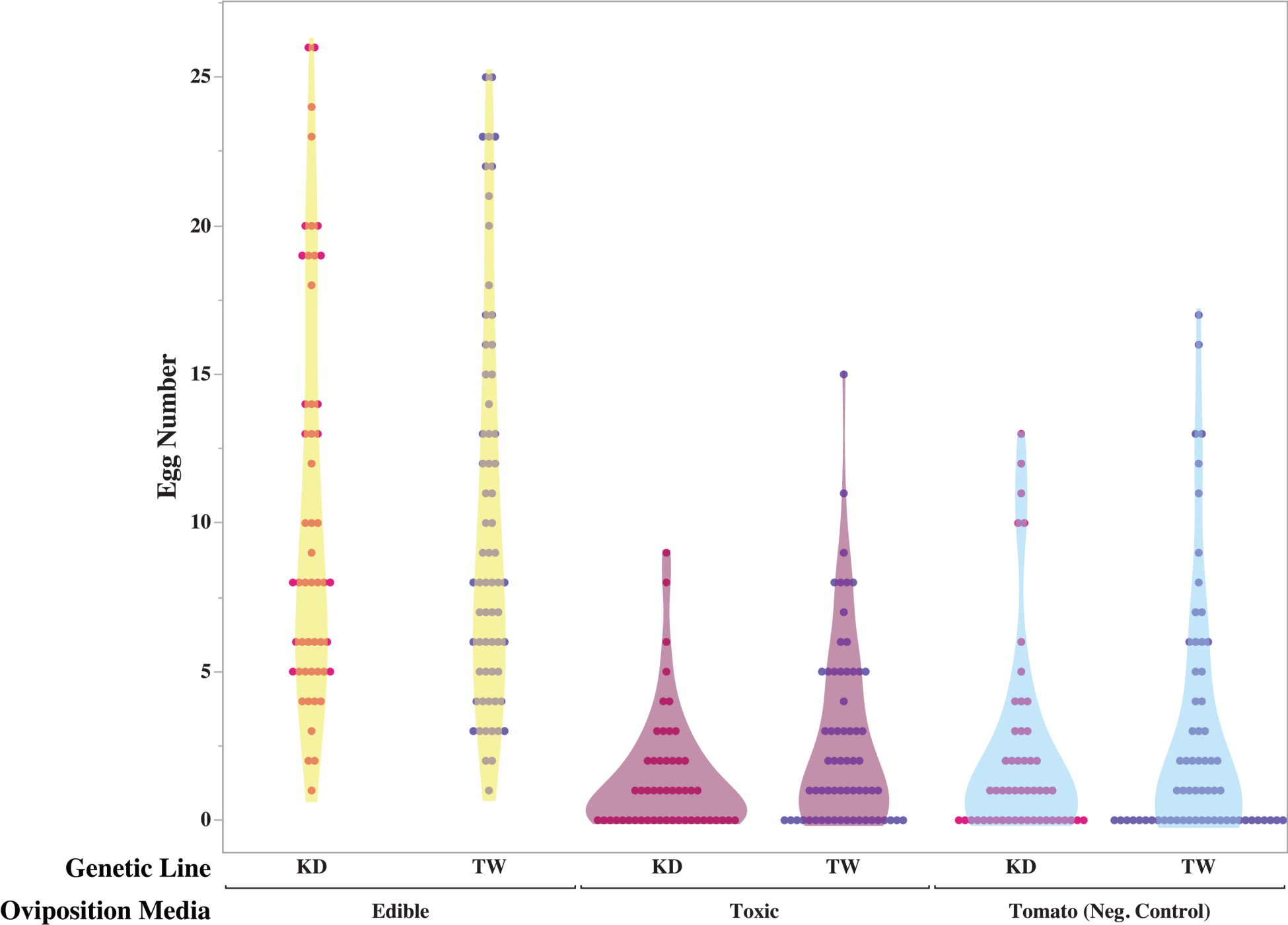
Visualization of the total number of eggs laid on each oviposition medium by the two lines across all replicates. X-axis indicates the media types.

**Table 2.** Comparison within each line of the mean number of eggs laid on the different media.

| Media | Line | Mean Eggs Laid | 95% CI |
| --- | --- | --- | --- |
| Edible Mushroom | KD<sup>§</sup> | 10.510*** | [9.251, 11.769] |
| Toxic Mushroom |  | 1.431 | [0.173, 2.690] |
| Tomato<sup>‡</sup> |  | 2.177 | [0.918, 3.435] |
| Edible Mushroom | TW<sup>†</sup> | 10.172*** | [8.965, 11.379] |
| Toxic Mushroom |  | 2.734 | [1.528, 3.941] |
| Tomato<sup>‡</sup> |  | 2.906 | [1.700, 4.113] |

\* $P < 0.05$ , \*\* $P < 0.01$ , \*\*\* $P < 0.001$
<sup>‡</sup>Negative Control
<sup>§</sup>51 replicates
<sup>†</sup>64 replicates

**Table 3.** Analysis of variance in oviposition preference.

| Phenotype | Source | DF | Sum of Squares | F-Ratio | Prob > F |
| --- | --- | --- | --- | --- | --- |
| Oviposition Preference | Line | 1 | 27.177 | 1.207 | 0.273 |
|  | Media | 2 | 4890.743 | 108.623 | <0.0001*** |
|  | Line x Media | 2 | 39.369 | 0.874 | 0.418 |

\*P < 0.05, \*\*P < 0.01, \*\*\*P < 0.001

### LARVAL PREFERENCE

During the completion of the oviposition preference assay, we observed that larvae would migrate from the medium where their eggs were laid (edible mushroom) to another (*e.g.*, toxic medium). To assess whether host preference occurs in larval *D. guttifera* of the KD line, eggs were laid on a plain agar medium with no nutritional value, and we measured the change in mass of the two mushroom based media (Figure 5). After 96 hours, the change in mass of the two media containing mushrooms (mean mass change > 1 gram for both media) was significantly higher (*P <* 0.001; Table 4) than for the plain medium (mean mass change = 646.53 mg) where the eggs were laid. When comparing the two, mushroom-based media, the change in mass did not differ significantly (*P >* 0.05). In characterizing the sources of variation in this data, we found that media type made the greatest contribution (*P <* 0.0001; Table 5). Additionally, the number of eggs laid on the plain medium was a significant source of variation (*P =* 0.0012). The interaction between media and the number of eggs was not significant (*P =* 0.563).

**Figure 5.**
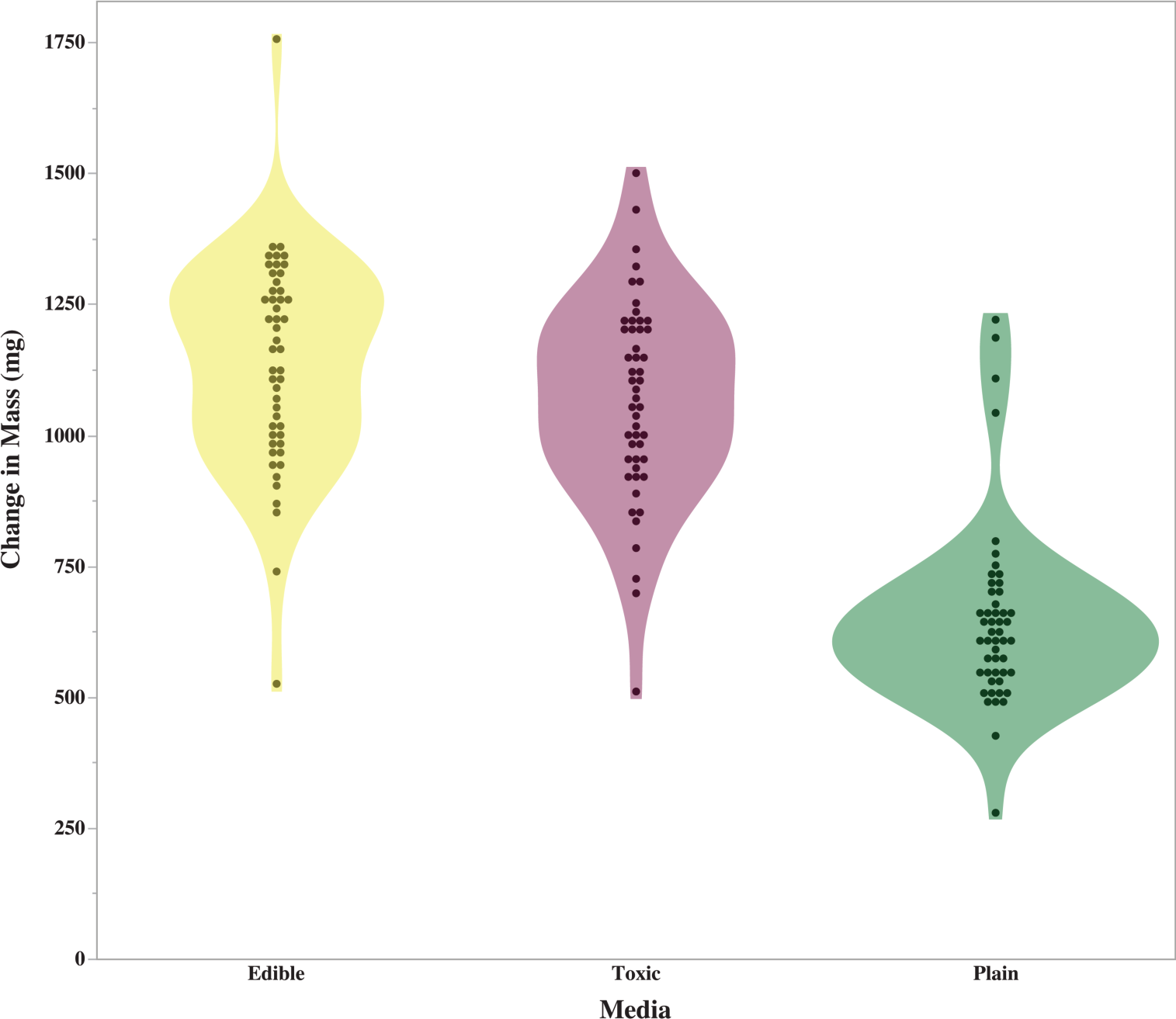
Comparison of the change in mass for each of the media types across the 50 replicates. X-axis indicates the type of medium.

**Table 4.** Comparison of mean change in mass of each media in the larval preference assay.

| Media Type<sup>§</sup> | Mean Change (mg) | 95% CI |
| --- | --- | --- |
| Edible Mushroom | 1135.06 | [1082.0, 1181.1] |
| Toxic Mushroom | 1067.61 | [1014.6, 1120.6] |
| Plain<sup>‡</sup> | 646.53*** | [593.5, 699.6] |

\*P < 0.05, \*\*P < 0.01, \*\*\*P < 0.001
<sup>‡</sup>Media eggs were laid on<sup>§</sup>50 replicates

**Table 5.** Analysis of variance of larval media preference.

| Phenotype | Source | DF | Sum of Squares | F-Ratio | Prob > F |
| --- | --- | --- | --- | --- | --- |
| Change in mass | Media | 2 | 7008512.3 | 103.405 | <0.0001*** |
|  | Egg # | 1 | 372547.8 | 10.9933 | 0.0012** |
|  | Media x Egg # | 2 | 39120.5 | 0.5772 | 0.563 |

\*P < 0.05, \*\*P < 0.01, \*\*\*P < 0.001

### COMPETITION

To characterize whether competition could alter oviposition preference in the *D. guttifera* KD line, we completed assays where the preferred oviposition site (edible mushroom medium) was either free from competition or already had eggs laid on it by either an inter- or intraspecific competitor. We counted the number of eggs laid on each medium and in close proximity (Figure 6). The mean number of eggs laid by the *D. guttifera* KD differed significantly between the inter- and intraspecific competition treatments (*P <* 0.05; Table 6). The flies laid fewer eggs (mean = 1.0714) when an interspecific competitor (*D. tripuncata*) had previously laid eggs on the edible mushroom medium. When considering the contribution of different factors in the data variation, competition was a significant variable (*P =* 0.0018; Table 7). Similar to the oviposition assay, the greatest source of variation in the number of eggs laid was media (*P <* 0.0001). However, there was also a significant interaction between media and competition (*P =* 0.0001).

**Figure 6.**
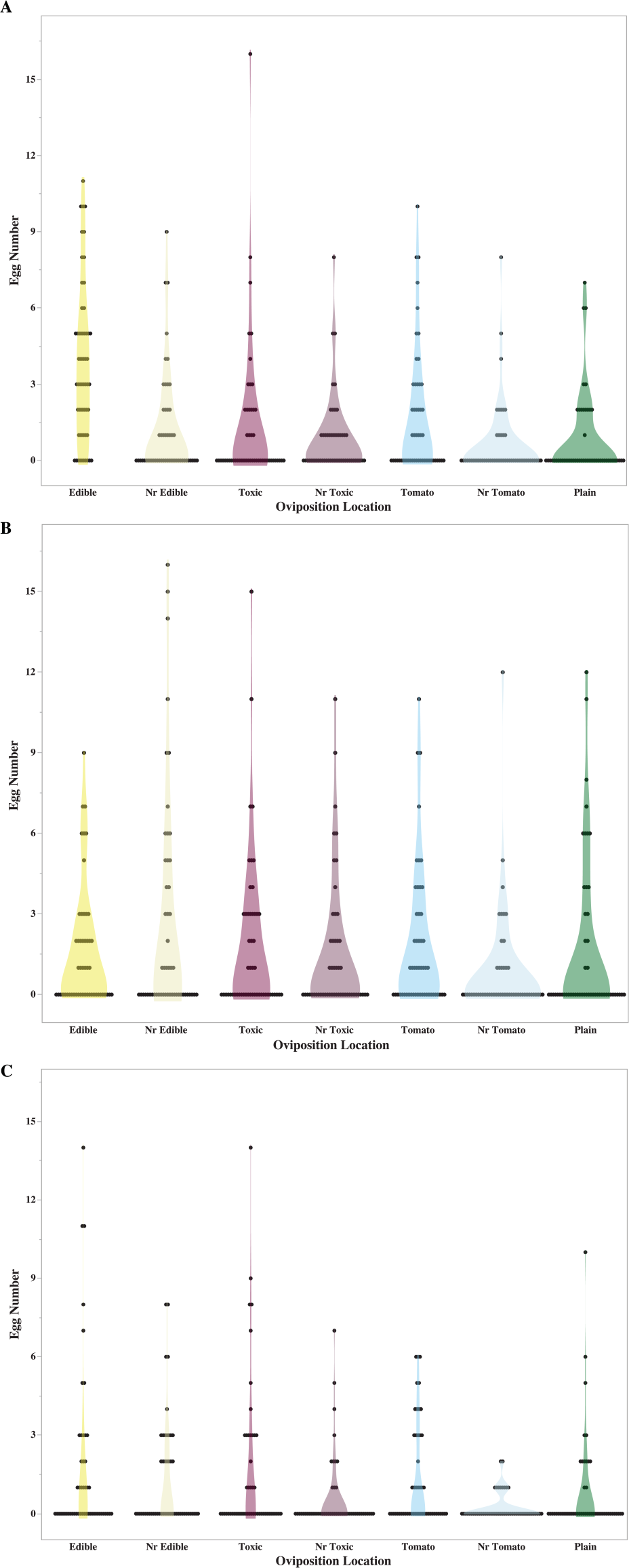
Assessment of the effect of competition on the number of eggs laid at each location. A) No competition, B) Intraspecific competition from *D. guttifera* TW line, C) Interspecific competition from *D. tripunctata*. X-axis indicates the different locations.

**Table 6.** Analysis of the impact of competition on the number of eggs laid.

| Competition<sup>§</sup> | Mean Eggs Laid<sup>‡</sup> | 95% CI |
| --- | --- | --- |
| Intraspecific | 1.706<sup>a</sup> | [1.446, 1.966] |
| Interspecific | 1.0714<sup>b</sup> | [0.812, 1.331] |
| None | 1.489<sup>ab</sup> | [1.229, 1.749] |

<sup>§</sup>52 replicates
<sup>‡</sup>Means not sharing any letters are significantly different in a Tukey-test at $P < 0.05$

**Table 7.** Analysis of variance of oviposition preference when experiencing competition and the impact on the mean number of eggs laid.

| Phenotype | Source | DF | Sum of Squares | F-Ratio | Prob > F |
| --- | --- | --- | --- | --- | --- |
| Oviposition Preference | Media | 6 | 333.621 | 9.339 | <0.0001*** |
|  | Competition | 2 | 75.738 | 6.361 | 0.0018** |
|  | Media x Competition | 12 | 230.698 | 3.229 | 0.0001*** |

\*P < 0.05, \*\*P < 0.01, \*\*\*P < 0.001

In our examination of the oviposition locations across the three treatments (Figure 6), we observed different patterns of host usage under each. However, the fewest eggs were laid in the plain media closest to the tomato across all assays (Table 8). The no competition treatment (Figure 6A) produced similar results to the original oviposition assay (Figure 4). The mean number of eggs laid was significantly higher on the edible mushroom medium (3.827, *P <* 0.05) in comparison to all other locations. In addition, the flies continued to lay more eggs on the tomato medium (mean = 1.731) than the toxic mushroom medium (mean = 1.346). When an intraspecific competitor laid eggs on the edible medium, the KD line laid most eggs near, but not on, the edible medium (Figure 6B). The mean number laid at this location was significantly higher (2.673, *P <* 0.05) than the near tomato location (0.827). The mean number of eggs laid at the other locations did not differ significantly from either of these two. Under the interspecific competition treatment (Figure 6C), the mean number of eggs laid in both the edible and toxic mushroom mediums (1.712 and 1.596 respectively) were significantly higher (*P <* 0.05) than the mean number laid near the tomato medium (0.231). Under both competition treatments, the number of eggs laid on the toxic mushroom medium was higher than the tomato medium.

**Table 8.** Examination of interactions between competition on media use in oviposition preference.

| Media | No Competition |  | Intraspecific (*D. guttifera*) |  | Interspecific (*D. tripunctata*) |  |
| --- | --- | --- | --- | --- | --- | --- |
|  | Mean Eggs Laid<sup>‡</sup> | 95% CI | Mean Eggs Laid<sup>‡</sup> | 95% CI | Mean Eggs Laid<sup>‡</sup> | 95% CI |
| Edible | 3.827<sup>a</sup> | [3.216, 4.438] | 1.596<sup>ab</sup> | [0.813, 2.379] | 1.712<sup>a</sup> | [1.126, 2.297] |
| Nr. Edible | 1.231<sup>b</sup> | [0.620, 1.841] | 2.673<sup>a</sup> | [1.890, 3.456] | 1.288<sup>ab</sup> | [0.703, 1.874] |
| Toxic | 1.346<sup>b</sup> | [0.736, 1.957] | 1.865<sup>ab</sup> | [1.0823, 2.649] | 1.596<sup>a</sup> | [1.011, 2.181] |
| Nr. Toxic | 0.865<sup>b</sup> | [0.255, 1.476] | 1.500<sup>ab</sup> | [0.717, 2.283] | 0.577<sup>ab</sup> | [-0.0082, 1.162] |
| Tomato (NC) | 1.731<sup>b</sup> | [1.120, 2.341] | 1.827<sup>ab</sup> | [1.0438, 2.610] | 1.308<sup>ab</sup> | [0.723, 1.893] |
| Nr Tomato | 0.615<sup>b</sup> | [0.0049, 1.226] | 0.827<sup>b</sup> | [0.0438, 1.610] | 0.231<sup>b</sup> | [-0.354, 0.816] |
| Plain | 0.808<sup>b</sup> | [0.197, 1.418] | 1.654<sup>ab</sup> | [0.871, 2.437] | 0.788<sup>ab</sup> | [0.203, 1.374] |

<sup>‡</sup>Means not sharing any letters are significantly different in a Tukey-test at $P < 0.05$

## DISCUSSION

A focal area in the study of plant-herbivore interactions is the evolutionary arms race between plants and/or fungi and the organisms that feed on them (Berenbaum and Zangerl 1998; Bernays and Graham 1988; Ehrlich and Raven 1964; Fraenkel 1959). Host defenses include structures that prevent/limit feeding damage, production of secondary metabolites to deter feeding, and phenological shifts (Agrawal et al. 2009; Hanley et al. 2007). The potency of the defensive chemicals produced by plants and fungi to deter feeding ranges from highly toxic to digestibility-reducing. The expectation of what organisms can feed on chemically defended hosts varies based on how noxious the compound is. Specialists evolve mechanisms that allow them to develop on hosts containing highly toxic compounds, but lose the ability to feed on a wide range of hosts (Cornell and Hawkins 2003; Whittaker and Feeny 1971). In turn, the detoxification mechanisms that are expected to evolve in generalists are those that will be useful against less toxic defenses that are common across multiple hosts (Ali and Agrawal 2012). We know far less about how novel adaptations to highly toxic compounds impact generalist species and how they are maintained. In this study, we examine this question in the mushroom-feeding *Drosophila guttifera*, which is broadly polyphagous on fleshy Basidiomycota (Sturtevant 1921), including toxic *Amanita* species. We used a combination of behavioral assays to assess how toxin tolerance impacts the life history of *D. guttifera* and the conditions that will drive this species to use a toxic mushroom host.

We first examined the occurrence of the novel adaptation of interest, cyclopeptide toxin tolerance, in the two available *D. guttifera* genotypes. Phylogenetic examinations of the evolution of this adaptation in the *quinaria* species group (Spicer and Jaenike 1996) and the *immigrans-tripunctata* radiation (Erlenbach et al. 2023) suggest that tolerance arose once and has been lost multiple times. In species that transition away from mushroom feeding, toxin tolerance is lost in ∼ 1 million years (Spicer and Jaenike 1996). As both *D. guttifera* genotypes have been maintained in the lab without exposure to cyclopeptide toxins for over a decade, we completed feeding assays where we reared larvae on diets without toxins and with either a single cyclopeptide (α-amanitin) or a natural toxin mix. The larvae of both lines survived on all diets; however, the patterns of survival varied between them. While treatment (presence or absence of cyclopeptide toxins) did not significantly affect survival of the KD line, it did significantly lower survival in the TW line. However, survival in both lines decreased on the diet containing the natural mixture of toxins. The lowered survival on the toxin mix could be due to synergistic and/or antagonistic interactions that occur within the mixture. Prior studies (Dyer et al. 2003; Richards et al. 2016) on tolerance of host secondary metabolites found that the potent bioactivity of some compounds is due to these types of interactions. When examining potential sources of the variation in the survival data across both lines, we found that there is a significant interaction between genotype and treatment (environment). This suggests that significant genetic variation is present for toxin tolerance in *D. guttifera*. With only two available genotypes, we are limited in the conclusions we can draw regarding genetic variation within the species. However, this finding is consistent with studies of cyclopeptide tolerance in other mushroom-feeding species within the *immigrans-tripunctata* radiation (Jaenike 1989; Kokate et al. 2022). Both studies identified a significant genotype by environment interaction for survival to adult when larvae are reared on diets containing cyclopeptide toxins. Kokate et al. (2022) identified this pattern in two other members of the *quinaria* group, *D. recens* and *D. falleni*. Thus, our results demonstrate that cyclopeptide tolerance is still present and is potentially a complex genetic trait.

Tolerance to highly toxic host compounds can allow organisms to exploit new niches and is associated with host specialization (Ehrlich and Raven 1964; Fraenkel 1959). Mushroom-feeding flies in the *immigrans-tripunctata* radiation offer the potential to observe the transition from generalist to specialist feeding strategies. To characterize a potential preference for the toxic host in female *D. guttifera*, we conducted oviposition assays using both genetic lines where female flies selected among edible and toxic mushrooms along with a negative control. If *D. guttifera* is transitioning to a specialized feeding strategy then we would expect that females exhibit a preference for cyclopeptide containing mushrooms. Such a pattern can be found in other *Drosophila* species, including *D. sechellia*, an ecological specialist on the chemically defended noni fruit of *Morinda citrifolia* (Álvarez-Ocaña et al. 2023). With our oviposition assays, we found that both lines exhibited similar preferences and laid significantly more eggs on the edible mushroom medium (*P <* 0.001). Somewhat surprisingly, both lines laid more eggs on the negative control (tomato) than the toxic mushroom medium. When examining the effect of different factors on oviposition variation, only media was significant. While these results would suggest that female flies show no preference for toxic mushrooms and are slightly more likely to lay eggs on a host that they do not use in the wild (tomatoes), work with other *Drosophila* species found that oviposition preference can be influenced by the proximity of the preferred substrate (Miller et al. 2011; Schwartz et al. 2012; Sumethasorn and Turner 2016). In these situations, female flies chose to lay eggs on a suboptimal substrate for larval development if the larvae could move to the optimal source. During the completion of the oviposition assays, we observed that larvae would migrate between the media in some replicates. Furthermore, this observed negative correlation between the female’s preference and a known host could be due to missing cues that would be present in a natural setting (Nylin et al. 1996; Thompson 1988). While it does not appear that a preference for toxic mushrooms is present in the female flies, there is the potential that our lab setting is missing ecological cues that drive usage of these hosts and thereby maintain cyclopeptide tolerance.

For polyphagous insect species, the addition of a new host source requires females to lay their eggs on the host as well as the larvae developing successfully. However, oviposition preference and larval performance are not always positively correlated (Gripenberg et al. 2010; Mayhew 1997; Murphy 2004). For species whose larvae are unable to move between distant hosts, such as *Drosophila*, it is counter-intuitive that females would exhibit a preference for laying their eggs on sub-optimal locations/hosts, but this strategy has been observed in *Drosophila* species when media/host options are close enough to allow movement (Schwartz et al. 2012; Yang et al. 2008). Given the absence of an oviposition preference for toxic mushrooms - and in fact a potential distaste for this host – as well as our observations that *D. guttifera* larvae moved among the oviposition media types, we conducted a feeding assay to assess whether larvae from the KD genotype showed a preference for edible or toxic mushrooms. To quantify preference, we assessed the change in mass of the two mushroom media and a plain agar medium after 96 hours. Studies of other *Drosophila* species found that larvae prefer specific yeast strains (da Cunha et al. 1951; Lindsay 1958) and that the stage of fungal decay impacts larval preference in other fungal-feeding species in the *immigrans-tripunctata* radiation (Kimura 1980). We found that both the type of media and the number of eggs present on the plain agar medium accounted for a significant portion of the variation. However, the interaction between these variables was not significant. When we compared the mean changes in mass of the media, we found that the larvae consumed significantly more (*P <* 0.001) of the media containing mushrooms than the plain medium. These results suggest that the larvae preferred food sources that contain necessary nutrients and that when more larvae are present more media is consumed, but the consumption of a specific type of mushroom medium is not influenced by the number of eggs. It does not appear that a preference for toxic mushrooms is present within the larvae. These results in combination with our findings from the oviposition assay raise the question of how cyclopeptide tolerance is being maintained in *D. guttifera*.

In a natural setting, the benefits of escaping from enemies and/or gaining access to a resource with fewer competitors can outweigh the negative costs associated with larvae developing on chemically defended hosts (Alzate et al. 2017; Craig et al. 2000; Murphy 2004; Singer et al. 2004). For mushroom-feeding *Drosophila*, Perlman et al. (2003) found that developing on a toxic mushroom reduced the load of *Howardula* nematodes, which can sterilize adults. While this suggests that the larvae of these species benefit from access to enemy-free space, adults that are infected with nematodes do not alter their behavior to seek out mushrooms containing cyclopeptide toxins (Debban and Dyer 2013). As such, we conducted additional oviposition assays to determine how intra- and interspecific competition affected host preference using the KD line of *D. guttifera*. In these assays, we provided the flies with the same oviposition media types, but the edible mushroom medium contained eggs laid by a different *D. guttifera* genotype or *D. tripunctata*, a mushroom-feeding species with overlapping distribution. In our no competition treatment, the results were consistent with what we had observed originally (*i.e.*, edible mushrooms strongly preferred over all other options). With the competition treatments, we found that the responses of the flies varied depending on the competitor. When eggs from an intraspecific competitor where present on the edible mushroom medium, the flies laid significantly more eggs (*P <* 0.05) than in the interspecific treatment. The preferred host medium varied between the two competition treatments (near edible – intraspecific and edible/toxic – interspecific). With both competitive treatments, the mean number of eggs laid on the toxic medium was greater than the tomato medium, unlike the oviposition assays without competition. Additionally, the variation in oviposition preference was significantly influenced by the interaction between media and type of competition. These findings are congruent with the work of Grimaldi and Jaenike (1984) who found that on edible mushrooms larvae experienced food shortages due to competition. In *D. recens*, another member of the *quinaria* group, higher levels of toxin tolerance were associated with lowered competitive ability (Kokate and Werner 2023). Our findings suggest competition and more specifically interspecific competition could play a role in the use of toxic host mushrooms. Thus, the ability to use resources with lower levels of competition could act to maintain cyclopeptide tolerance in *D. guttifera*.

Overall, the results of our study do not provide any support for the hypothesis that *D. guttifera* is transitioning to a specialized feeding strategy on toxic mushrooms. Moreover, while cyclopeptide tolerance is present in both lines, the adults show no preference for toxic mushrooms, using them at rates equivalent to non-natural hosts (tomatoes). Given how rapidly this adaptation can be lost (∼1 million years; Spicer and Jaenike 1996), we assessed the impact of intra- and interspecific competition on toxin tolerance maintenance. Our results suggest that competition can cause flies to make use toxic host mushrooms. Further studies could investigate the role of parasitoid wasps and the ephemeral nature of mushrooms in host choice in this species. Overall, our assays provide further evidence for the role of competition in driving the selection of hosts with associated costs in larval fitness.

## Supporting information

Impact of treatment on larval performance in both D. guttifera genotypes

## Acknowledgments

This work was funded by grants from the National Science Foundation (DEB-1737869 to CSC and Laura Reed – University of Alabama and DBI-2217912 to CSC) and startup funds provided by Appalachian State University to CSC.

## DATA AND CODE AVAILABILITY

Data and JMP code have been deposited in the Dryad Digital Repository

(https://doi.org/10.5061/dryad.98sf7m0pm) and Zenodo

(https://doi.org/10.5281/zenodo.8280572)

